# Robust statistical assessment of Oncogenotype to Organotropism translation in xenografted zebrafish

**DOI:** 10.1101/2025.05.28.656734

**Authors:** David Saucier, Xuexia Jiang, Divya Rajendran, Roshan Ravishankar, Erin Butler, Aruna Marchetto, Raushan T. Kurmasheva, Thomas G.P. Grünewald, James F. Amatruda, Gaudenz Danuser

## Abstract

Organotropism results from the functional versatility of metastatic cancer cells to survive and proliferate in diverse microenvironments. This adaptivity can originate in clonal variation of the spreading tumor and is often empowered by epigenetic and molecular reprogramming of cell regulatory circuits. Related to organotropic colonization of metastatic sites are environmentally-sensitive, differential responses of cancer cells to therapeutic attack. Accordingly, understanding the organotropic profile of a cancer and probing the underlying driver mechanisms are of high clinical importance. However, determining systematically the organotropism of one cancer versus the organotropism of another cancer, potentially with the granularity of comparing the same cancer type between patients or tracking the evolution of a cancer in a single patient for the purpose of personalized treatment, has remained very challenging. It requires a host organism that allows observation of the spreading pattern over relatively short experimental times. Moreover, organotropic patterns often tend to be statistically weak and superimposed by experimental variation. Thus, an assay for organotropism must give access to statistical powers that can separate ‘meaningful heterogeneity’, i.e., heterogeneity that determines organotropism, from ‘meaningless heterogeneity’, i.e., heterogeneity that causes experimental noise. Here we describe an experimental workflow that leverages the physiological properties of zebrafish larvae for an imaging-based assessment of organotropic patterns over a time-frame of 3 days. The workflow incorporates computer vision pipelines to automatically integrate the stochastic spreading behavior of a particular cancer xenograft in tens to hundreds of larvae allowing subtle trends in the colonization of particular organs to emerge above random cell depositions throughout the host organism. We validate our approach with positive control experiments comparing the spreading patterns of a metastatic sarcoma against non-transformed fibroblasts and the spreading patterns of two melanoma cell lines with previously established differences in metastatic propensity. We then show that integration of the spreading pattern of xenografts in 40 – 50 larvae is necessary and sufficient to generate a *Fish Metastatic Atlas* page that is representative of the organotropism of a particular oncogenotype and experimental condition. Finally, we apply the power of this assay to determine the function of the *EWSR1::FLI1* fusion oncogene and its transcriptional target SOX6 as plasticity factors that enhance the adaptive capacity of metastatic Ewing sarcoma.

## Introduction

The ability of cancer cells to grow under adverse conditions is a defining feature of the transformed phenotype, and the primary driver of adverse outcomes in people with cancer. To survive and proliferate, cancer cells must overcome not only cell-autonomous barriers, such as cell-cycle checkpoints and apoptosis, but also external stresses including limited availability of nutrients and oxygen, attack by immune cells, diverse tissue microenvironments, and the effects of chemotherapy and radiation therapy. Thus, cancer is a disease of cell adaptation to adverse conditions [1, 2]. Biologically, this adaptation manifests itself in, and is likely enabled by, heterogeneous behaviors at the single cell level, as cancers cells with diverse phenotypes exhibit differential ability to adapt to diverse environments.

In the context of a whole organism, this adaptation is directly related to organotropism, i.e., the propensity for tumor cells to spread to different tissues and organs during the process of tumor dissemination and metastasis. Clinically, organotropism is a critical issue, as it strongly impacts treatment success and survival of patients with cancer, because tumor masses present in different organs can exhibit differential responses to therapy. This is true for a wide spectrum of both adult and pediatric malignancies, including epithelial cancers such as breast [3, 4], colorectal [5] and urothelial carcinoma [6], for testicular germ cell tumors [7] and for sarcomas including Ewing sarcoma [8, 9] and osteosarcoma [10, 11]. The organotropism exhibited by different cancers, and the differential response of tumor cells in different tissue environments to therapy, most likely reflect the dynamic interplay between phenotypically heterogeneous cancer cells and the different tissue microenvironments encountered by those cells. However, the specific mechanisms that determine this interplay are poorly understood.

Advanced genomics techniques including next-generation and single-cell sequencing have revealed the presence of genetic subpopulations in tumors as one source of such heterogeneity, lending support to models in which different environmental conditions exert pressures resulting in selective survival of specific subclones [12–17]. Indeed, for some tumor types there is evidence that genetic subclones exist at the time of diagnosis, and undergo expansion as a consequence of therapy [18–20]. Other studies, however, have found similar representation of driver mutations in primary and matched metastatic lesions, with little evidence that somatic genomic variation enables cell adaptation [21–23]. While these findings may be due in part to the difficulty of detecting rare variant subclones within a primary tumor, there is also evidence to suggest that cancer cells may adapt dynamically over relatively short time scales. This acute adaptation to environmental conditions can occur at the level of signaling, allowing cells to alter their metabolism or morphology, undergo epigenomic reprogramming, or enter quiescent states in response to the stresses exerted by particular environments [21, 24–29]. Just as with genomic heterogeneity, this type of dynamic non-genomic heterogeneity can modulate transcriptional, translational, and posttranslational programs, ultimately shaping key functions for cellular homeostasis and proliferation. These mechanisms may be especially relevant for pediatric cancers such as Ewing sarcoma, in which tumor cells may exhibit only a single somatic mutation [30, 31]. However, comparatively little is known about these mechanisms of adaptation, or to what extent different cancers with very different mutation spectrum and mutation burden may share common adaptive mechanisms.

To make progress on this problem, new tools are needed to efficiently screen variation in organotropism between cancers with different genetic and molecular makeup. To be most powerful, these tools would ideally be combined with assays that permit monitoring of the process of selection and acute adaptation in situ. The introduction of human cancer cells as xenografts into immunocompromised mice is a powerful approach, but poses challenges including the difficulty of performing live imaging at rapid timescales, as well as logistical difficulties and expense that can limit the number host organisms and thus the number of observable events [32–36]. As a complement to mice, zebrafish larvae have several key advantages for the assessment of heterogeneous cell phenotypes in diverse environments. Zebrafish oocytes are fertilized external to the mother fish, and 48 hours after fertilization the optically transparent, 1 mm-long larva has fully functional organs with the same spectrum of tissue mechanics (soft, stiff, dense, porous, intravascular) as human tissues [37–42]. Human cancers readily engraft into zebrafish, directly interact with the zebrafish microenvironment, including immune cells [43, 44], and exhibit angiogenic [45] and metastatic [45, 46] behavior.

The central task at hand in measuring organotropism of tumor cells is separating ‘meaningful heterogeneity’, i.e., heterogeneity that determines organotropism, from ‘meaningless heterogeneity’, or heterogeneity that causes experimental noise. This is challenging as the meaningful heterogeneity may generate weak patterns, and thus the signal to noise ratio may be low. The key question we set out to address is whether large numbers of experimental repeats allow these patterns to emerge from the noise of meaningless heterogeneity. The features of the zebrafish system—large clutch size, optical transparency, rapid development and the presence of a diverse range of vertebrate organ—make it an ideal experimental model for this purpose. Here we introduce Fish Metastatic Atlas (FMA) as a means to uncover meaningful heterogeneity in the metastatic spread of human cancer xenografts. The assay leverages the medium throughput imaging of entire zebrafish larva to integrate the spreading patterns of human cancer xenotransplants into a consensus distribution of organotropism for a particular oncogenotype. The readout of organotropic behavior relies on the measurement of cancer cell locations, i.e. the assay captures a mix of bona fide metastatic colonies, random cell spreading, and cells trapped in the vasculature. For an individual larva these events are indistinguishable. However, our study shows that by combining the location patterns of 40 – 50 the meaningful heterogeneity associated with the systematic visitation of specific tissue niches outsizes the meaningless heterogeneity associated with random events. We further equip our pipeline with statistical metrics to compare consensus distributions of distinct oncogenotypes. Importantly, even though the assay detects with high sensitivity shifts in the organotropism of different conditions, the origin of the shifts, i.e. differential behavior along the metastatic cascade, cannot be determined. To acquire this information, the proposed assay must be combined with cell-level analyses of metastatic functions (migration, immune evasion, survival, and proliferation).

## Results

### Harnessing zebrafish embryos as host organisms for human cancer xenotransplantations

To harness the power of the zebrafish model system for the study of organotropism we performed xenotransplantations of human cancer cells into the perivitelline space of anesthetized zebrafish larvae. Specifically, we injected a labeled human cancer cell suspension in larvae at 2 days post fertilization using a pulled-capillary needle and micro-injector (Fig S1A). Per session, injections were repeated for 100-180 fish resulting in 90-120 larvae containing fluorescently labeled human cancer cells. Between 10 and 25% of the successfully injected fish display cells in the vasculature immediately after injection. These larvae were removed from the petri dish as the ensuing distribution of viable cancer cell colonies off the injection would not be representative of a bona fide metastatic spreading process subject to invasion and intravasation. Of the remaining larvae, 15-30% do not survive the first 48 hours post injection (Fig S1B). Hence, the typical harvest of larvae suitable for imaging amounts to ∼50 samples per injection session. For a particular cancer cell type, we conducted 3-5 experimental runs, resulting in 150 – 250 imageable larvae.

For the analysis of patterns of organotropism larvae were fixed by paraformaldehyde and prepared for imaging 48 hours after injection. Image acquisition was performed in four channels to determine autofluorescence (green emission), cell signal (red emission), fish morphology (brightfield), and a fish outline estimate via a Leica LasX generated depth label map (see Methods). Stacks of 12 – 20 z-positions were acquired in steps of 42 – 50 microns satisfying the requirements of Nyquist sampling given a depth of field between 85 and 100 microns dependent on the zoom setting (see Methods). Images of the individual z-positions were then merged into a full focus projection image, in which each pixel is labeled by the image layer that contributes locally to the in-focus information. Depth label maps were more reliable than the brightfield channel for segmenting regions of low contrast such as the fin region, a site of interest for studies in cancer cell invasion.

### Registration individual fish larvae to a reference template

The experimental success rate would drop precipitously if fish with non-stereotypical morphologies were excluded. Only 20-30% of the larvae display the canonical morphology with straight dorsal alignments. Given the drop out due to cell deposition in the vasculature and larval perishing, a selection of only the larvae with canonical morphology would reduce the experimental harvest to less than 10% of the original spread, insufficient for an assay targeting the identification of rare but systematic metastatic deposition events. Thus, we sought to maximize the harvest in each session by aligning non-ideal fish with a canonical template fish morphology.

Determining systematic differences in organotropic patterns between oncogenic conditions is complicated both by the vast heterogeneity of organ access by cancer cells with identical genomes between individual zebrafish larvae and by the diversity of embryo morphology (Fig. 1A). The decision whether two oncogenic conditions generate statistically distinct or identical organotropic patterns (Fig. 1B) requires the definition of a robust fish larval reference system for accumulation of a sufficiently large number of metastatic spreading instantiations in different larvae such that a stereotypical organotropic pattern for an oncogenotype or experimental condition can emerge over random experimental variation. This fish larval reference system will also enable the statistical comparison of accumulator patterns between experiments.

**Fig. 1.**
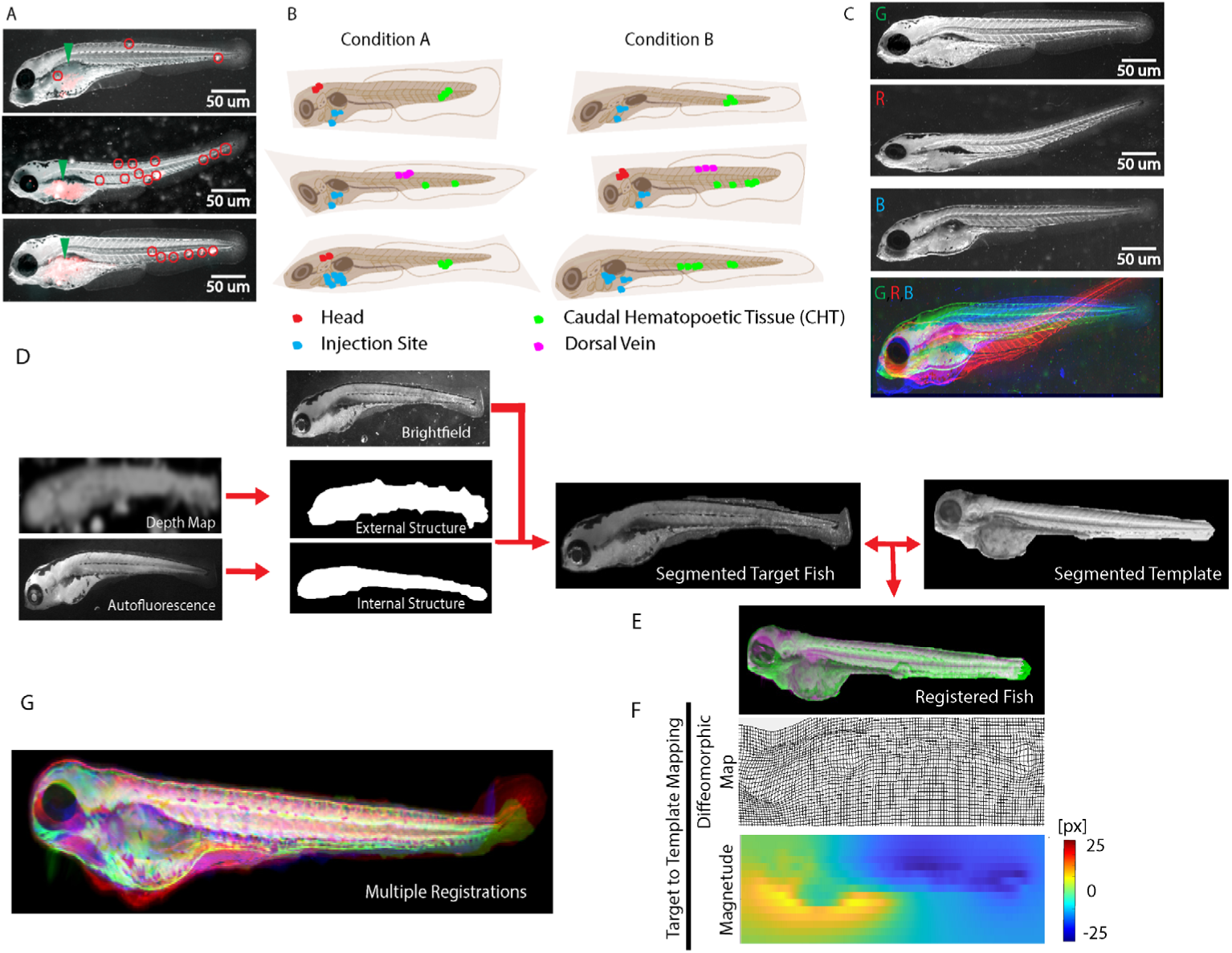
Definition of a fish larval reference system to assess shifts in organotropism between human cancer xenografts. **(A)** Three representative examples of zebrafish larvae (gray brightfield images) injected with 90 – 120 TC32 human Ewing sarcoma (ES) cells labelled with cytosolic mRuby reporter (red colors overlaid). Injection sites are indicated by green arrowheads. The zebrafish display different gross morphologies. Cell colonies form in highly variable locations (red circles). **(B)** Cartoon defining some of the zebrafish gross anatomy. Given the variation on gross morphology and colony formation, whether organotropism under experimental Condition A differs from Condition B cannot be directly addressed. **(C)** Overlay of three injected larvae in RGB pseudocolors registered by center of mass alignment. **(D)** Registration pipeline by diffeomorphic maps. Inputs are the brightfield and autofluorescence (captured in the band 440-470 nm) images plus depth map reflecting the 2.5D topography of the larva. The external and internal outlines of the larvae are extracted by Rosin thresholding of the depth map and active contour segmentation of the autofluorescence signal, respectively. Segmented images of target fish larvae are registered to a segmented image of a template fish larva via diffeomorphic mapping of respective brightfield images. **(E)** Registration is visualized by green/magenta overlays of the two brightfield images. **(F)** The diffeomorphic mappings represented by grid displacements from target to template (top) and by magnitudes of the displacements (bottom). **(G)** RGB overlay of the three embryos shown in (C) after registration. Mismatches are visible in the areas of the fins, especially in the tail region, whereas the brightfield signals of the bodies show good conservation of morphology after registration.

Although zebrafish at 48 hours post-fertilization display a highly stereotypical gross anatomy [37], details in the development of the animal morphology as well as variation in the sample mounting result in significant geometric mismatch between individual larvae (Fig. 1C, Video 1). The majority of previous studies of metastatic spreading in zebrafish have not attempted to analyze specific patterns of secondary tissue colonization distribution but only implemented metrics of the overall spreading that were relatively agnostic to the gross-morphological variation between samples [47–52]. The few studies that do compare organ and tissue occupancy approached the definition of a common coordinate framework across multiple specimens by selection and center-of-mass alignment of straight-dorsal embryos [53–55]. Similarly, studies of metastatic patterns in adult, immunocompromised zebrafish relied on a direct overlap of specimens with similar shapes followed by generation of heatmaps of tumor clusters [56–59]. However, reliance on a pre-selection of a typically small subgroup of geometrically suitable fish is tedious and lacks the reproducibility required for robust statistical evaluation of organotropic patterns. Even with larvae of fairly consistent gross-morphology it is evident that an alignment by translocation and rotation, and perhaps also scaling, prohibits the identification of specific host tissue environments across even a small cohort of fish (Fig. 1C). We therefore set out to implement an alignment procedure that establishes a diffeomorphism between individual specimens in a cohort of fish and a template fish serving as a reference.

We leveraged the recent development of a registration algorithm [60] that preserves diffeomorphism, i.e. a bidirectional mapping, between fine-grained patterns to register the brightfield image of a target fish to the brightfield image of a template fish (Fig 1D). To increase the precision of the registration the algorithm makes use of the contours of the segmented target and template fish. These contours are often ill-defined in the brightfield and autofluorescence channels, especially around the thin and weakly structured fins. To ameliorate the outline of this poorly defined outer boundary we applied an active contour segmentation to the depth label map, initialized by the conservative estimate of the perimeter of the embryo visible in the autofluorescence channel. The registration algorithm defines the diffeomorphic pixel-by-pixel mapping between target and template by concurrently maximizing the alignment of the two fish boundaries and texture overlap in the interior, subject to the constraints that i) each finite area in the target fish image corresponds uniquely to a finite area in template fish image and ii) the topology of finite areas is preserved. Contrary to most image registration algorithms, which meet the requirement of topology preservation by applying smoothing filters to eliminate potentially confusing fine-grained image structures, our implementation preserves topology in an iterative sorting of mapping vectors. This allows the registration of small but salient image features [60], such as those generated by individual organelles. The effect and quality of diffeomorphic mapping can also be visualized by color-overlay between registered target fish and template (Fig 1E). The diffeomorphism between target and template fish image can be visualized by the deformation of a grid, which indicates the displacement of grid points in the target image necessary to match up with the corresponding location in the template fish image (Fig. 1F, top). Of note, despite the locally strong grid deformation, none of the grid lines cross, which validates the preservation of topology. The displacement magnitude can be displayed in a color map (Fig. 1F, top). Application of the same pairwise mapping to a series of target fish images registered to the same template results in the alignment of an entire experiment in a common coordinate framework (Fig. 1G).

### Elimination of larvae with gross registration defects

The objective of our pipeline is to accumulate data from a collection of fish larval samples that is large enough to draw meaningful conclusions on putative differences in the distribution of metastatic events between xenografts with distinct genetic backgrounds or between other experimental variation. The distribution differences can be relatively subtle and concentrated, for example, in a single location. Hence the registration process had to be equipped with tools to automatically assess the quality of alignment and eliminate outliers that would deteriorate the accuracy of the integrated distributions. Fig. 2A presents a gallery of brightfield images of 30 injected and fixed larvae, randomly pulled from a collection of >1,000 larvae. This unbiased view indicates the significant variation in fish gross-morphology as well as imaging conditions (pitch angle changes, particles and debris in the background, etc.) that complicate the registration of larvae collections to a common template. We have implemented a two-step filtering process to eliminate gross misalignments that cannot be addressed by diffeomorphic mapping. Fig. 2B shows an example of two target fish images that fail (Target 1) or succeed (Target 2) in passing the first step, which filters out targets with exceedingly poor overlap with the template. Each target in an experimental series is translocated so that its mask representing the external structure has maximal overlap with the mask of the template (Fig. 2C). Subsequently, we assemble a histogram of the fractional overlap (Fig. 2D) and reject the 10% lowest fractional overlap targets. The targets that pass the filter are then aligned to the template by diffeomorphic registration. Target fish images with average displacement magnitudes well outside (Fig. 2E, top) the normal range of displacements (Fig. 2E, bottom) constitute a potentially poor registration due to local mismatches in fish morphology that cannot be captured by a global diffeomorphism. Similar to the first filter step, 10% of the targets with the largest displacements are rejected (Fig. 2F). Application of the combined filters to the 30 fish in Fig. 2A demonstrates the robust outlier selection by this relatively simple strategy (Fig. 2G).

**Fig. 2.**
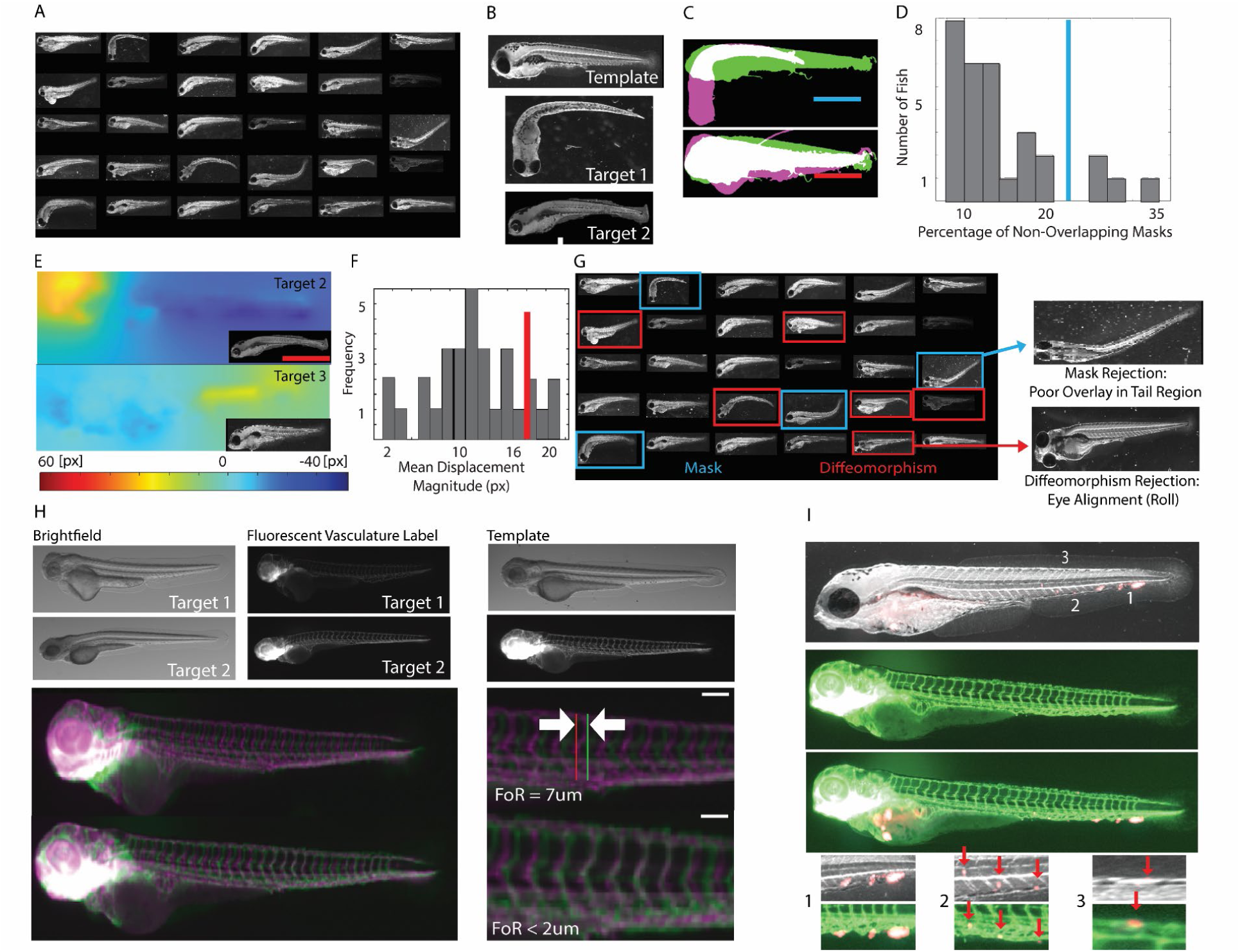
Quality assessment and filtering of gross alignment errors during registration. **(A)** Random selection of 30 fish larvae from a database of N>1000 specimens indicating significant differences in gross-morphology and imaging orientations. **(B)** Brightfield images of a template and two targets (target 1 with obvious tail bend and roll of the entire corps; target 2 with relatively minor morphological deviation from the template. **(C)** Mask overlays of target 1 (magenta) to template (green, top) and target 2 (magenta) to template (green, bottom) after coarse translocation alignment. **(D)** Histogram of the percentage of non-matching mask area (colored area in C) in relation to the mask union for N = 30 target-template pairs. 10% of targets with the highest mismatch (blue line) are eliminated. **(E)** Magnitude of displacements for target 2 (top) and target (3) (insets, bright field images) in diffeomorphisms to the template shown in (B). **(F)** Histogram of mean displacement magnitude in diffeomorphisms for N = 30 target-template pairs. 10% of targets with the highest displacement magnitude (red line) are eliminated. **(G)** Overview of embryos removed from the selection in (A) based on mask mismatch (blue boxes) and diffeomorphism displacements (red boxes). One example removal per filter criterion indicates that a major cause of data elimination is rotation of the body or head of the embryo in the dish. **(H)** Assessment of registration quality based on vasculature. Images (brightfield and fluorescent vasculature reporter Tg(*fli:gfp*) are shown for two targets and a template embryo. Registration performed using the brightfield, depth map (not shown) and autofluorescence (not shown) information. Using the determined diffeomorphisms for registration the deformed vasculature of the target embryos (shown in magenta) is overlaid to the vasculature of the template embryo (shown in green). Both examples show a registration match of 2-7 microns. **(I)** Colonies of fluorescent cells in a target (top) are mapped into the vasculature image of a template fish (middle and bottom). Three example colonies are located in the caudal hematopoetic tissue (1), intersomitic artery (2), and dorsal longitudinal anastomotic vessel (DLAV) (3), indicating precise alignment between target and template.

### Validation of the registration pipeline

To test the performance of the registration pipeline we used a genetic reporter, Tg(*fli:gfp*), to label all vasculature and endothelial regions of the zebrafish larvae at the time of injection (2dpf). Of particular interest is the intersomitic vasculature along the trunk of the fish flanked by the dorsal longitudinal anastomotic vessel (DLAV) and caudal vein. These periodic, stereotypical structures are conserved across larvae and are an excellent metric for registration quality.

We established diffeomorphisms between two target fish and a template based on only the brightfield image data. We then applied the diffeomorphisms to project the vasculature of the target onto the vasculature of the template (Fig. 2H). We defined the Fidelity of Registration (FoR) as the distance from the midline (15^th^) intersomitic artery between the two registrations. The two examples yielded representative FoR values of ∼7 and ∼2 microns, respectively. Of note, this registration mismatch is significantly smaller than the typical diameter of a metastatic colony of 20 – 30 microns. Hence, the registration is adequate to bring metastatic deposits associated with the same organ into overlap in the template space, allowing the statistical evaluation of the frequency of visitation of a particular microenvironment across many fish.

Finally, we wondered whether we could use the registration to align observations of metastatic colonies imaged in one fish with observations of a reference tissue like the vasculature imaged in another fish (Fig. 2I). Three sample regions indicate highly plausible positioning of colonies in between vessels as well as at vessel walls, affirming that the majority of metastatic spread visible throughout the fish body after injection associate with nascent metastatic colonies seeded from extravasating cells.

### Accumulation of individual metastatic spreading patterns in the coordinate frame of the reference template

Equipped with a mechanism to adjust variation in gross morphology between individual fish embryos, we could proceed to testing whether the metastatic spread of xenografted cells of a particular oncogenotype would follow an organotropic pattern. We hoped to accomplish this by overlaying the fluorescent channels diffeomorphically aligned to a template fish (Fig. 3A). Under significant molecular control of organotropism, the summation of these channels in an accumulator would be expected to yield a fluorescent pattern with marked hotspots. Although few hotspots were discernible, especially in the tail (see Fig. 3A, arrows), the accumulator signal overall was dominated by autofluorescence. This unveiled a notorious challenge in detecting organotropic patterns. Even if a few cells tend to systematically colonize a particular region of the fish, the variation in the visitation between fish is high. Moreover, at an observation point 48 hours post injection, colonies in individual fish may consist of 1 – 5 cells. In the majority of anatomic regions in the fish, the fluorescent intensity of such colonies may exceed the autofluorescence background by 2 – 3 times at best. Hence, a location that that is visited in every other or every third fish will not appear as a fluorescent signal hotspot in a simple summation accumulator.

**Fig. 3.**
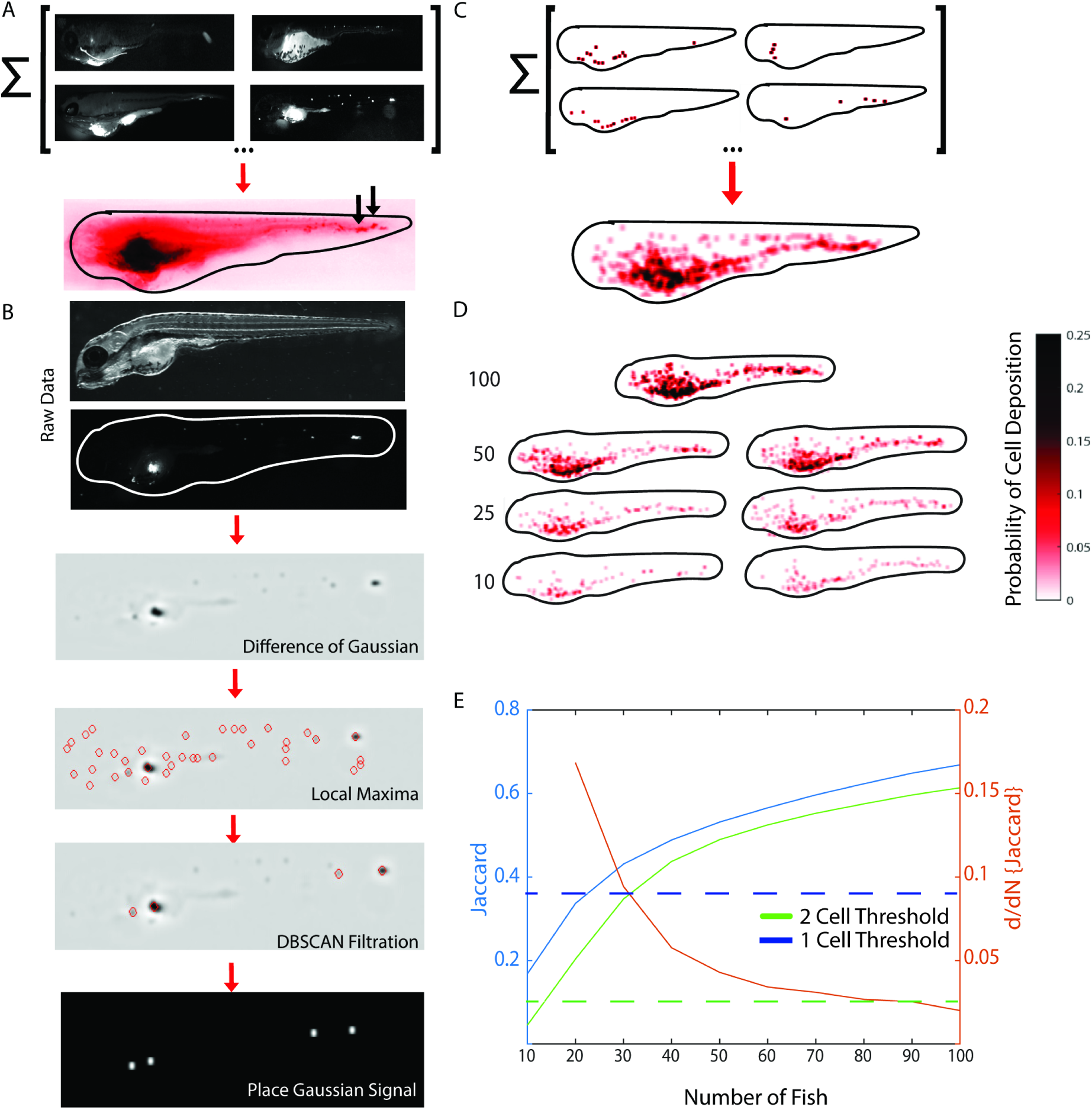
Assembly of accumulators representing the organotropism for a particular oncogenotype. **(A)** Sum of fluorescent images of injected cells is dominated by autofluorescence signal. Shown are four individual embryo images registered to a common template contributing to an accumulator of N = 30 embryos. With the exception of a few bright colonies in the dorsal part of the tail (arrows), the accumulator presents a diffuse signal stemming from the systematic addition of an autofluorescence signal in every contributing embryo. **(B)** Image processing pipeline to suppress the effect of autofluorescence. Brightfield image of an embryo injected with fluorescent cells. The fluorescence channel is filtered with a difference of Gaussian (DoG) kernel to amplify punctate signals above diffuse background, followed by local maxima detection and filtration by density-based spatial clustering (DBSCAN). The remaining points indicate loci of high probability of cell colony formation. These points are copied into a particle-based accumulator as a normalized Gaussian signal with a pre-calibrated width corresponding to the average image width of an isolated colony. **(C)** Accumulator of particles including the same embryos as shown in (A). **(D)** Accumulators of N = 100 embryos and of pairwise randomized sub-cohorts of N = 50, 25, and 10 embryos. The accumulators are normalized to report the local probability of a cell deposition. **(E)** Reproducibility of accumulators as a function of the size of the sampling cohort, assessed by the Jaccard index between accumulator pairs randomly sampled from the full data set of N = 100 embryos. Blue and green curves, evolution of Jaccard indices using the minimum of 1 and 2 cell deposits as a criterion for accumulator binarization. Each curve represents the mean of r = 2000 accumulator pairings. Dashed blue and green lines, Jaccard index for accumulator pairs with randomized cell deposits. Red curve, derivative of the Jaccard index with respect to sampling cohort size (using the t = 2 cell deposit Jaccard index curve).

To overcome the odds of losing sporadic but systematic colonization events to autofluorescence we implemented a particle-based accumulator (Fig. 3B). We applied Difference-of-Gaussian (DOG) filtering and local maximum detection to suppress local background around potential colonies. The resulting selection of particles contained a large number of maxima associated with noise. Moreover, colonies with multiple cells slightly spread apart generated several maxima in close proximity, i.e. the magnitude of the maxima may be quite low despite an overall relatively strong fluorescent signal for the colony. Under these circumstances particles representing bona fide colonies cannot be selected by straightforward thresholding. Instead, we applied a density-based spatial clustering (DBSCAN), which accounts for the local particle density as well as intensity to identify meaningful cluster centers. Finally, to integrate the selected colony particle fields into an accumulator (Fig. 3C), we centered a unit Gaussian signal at the location of each DBSCAN detection. The width of the Gaussian was chosen to reflect the average pixel coverage of a single colony image. Of note, by applying a unit Gaussian and after division of the summed particle fields from individual fish by the total number of individual fish, the accumulator is readily interpretable as a probability map for metastatic colonization: The accumulator value *A*(x,y) at any location inside the template fish outline indicates the probability of finding a colony at the diffeomorphically matching location in any of the individual fish.

### Interpreting accumulator patterns

Two questions arise with this accumulator strategy: 1) For a particular oncogenotype, is there a characteristic accumulator pattern, or is the accumulator merely representing a random spread of cells? 2) How many fish are necessary to capture a characteristic pattern? To address this, we performed a power analysis using a data set of 100 fish injected with the TC32 Ewing sarcoma (EwS) cell line. EwS is a malignant bone and soft-tissue cancer that occurs most commonly in children, adolescents, and young adults and is most commonly caused by a t(11;22) chromosomal translocation that creates the *EWSR1::FLI1* fusion oncogene [30, 31]. We bootstrapped accumulators from N fish randomly selected from the total of 100, permitting repeats [61, 62].

Visually, the accumulator patterns seem to stabilize at N ∼ 50 fish (Fig. 3D). Below N = 50 fish, the colonization patterns in ventral tail and dorsal vasculature could not be reproduced. To quantify the pattern stability, we computed for a given N 5000 bootstrapped accumulator pairs and compared their colonization patterns using the Jaccard coefficient, i.e. the ratio of intersection to union of the paired patterns. To compute the Jaccard coefficient accumulator patterns were binarized at a threshold of 1/N or 2/N, i.e. we only accept pixels that contain at least one or two visitations, respectively, in one of the contributing individual fish. As expected, the Jaccard coefficient monotonically increases to values of 0.67 and 0.61, respectively, at N = 100 fish, indicating significant randomness even at the full data set. These asymptotic coefficient values compare to baseline Jaccard coefficients for complete randomization of the accumulator patterns of 0.37 (1/N) and 0.10 (2/N), respectively. The 1/N and 2/N curves cross the baseline values at 22 and 14 fish. This means that any analysis of metastatic spread with less than ∼20 fish is not distinct from a random pattern analysis. Beyond these minimal criteria for fish number the slope of the coefficient curves monotonically decreased, which confirms convergence of the patterns. For N ∼ 45 fish the change in Jaccard coefficient compared to the coefficient computed for N – 10 fish dropped below 5%. This point also coincides well with the point of inflection in the curve of convergence, i.e. beyond 45 fish the gain in reproducibility of the accumulator pattern is fairly insignificant, matching with the visual impression in Fig. 3D. Therefore, for most conditions investigated in this study we applied the criterion of 45 fish as the requirement for characterizing the metastatic pattern of an oncogenotype. Moving forward we refer to one such characterization as a page of a *Fish Metastasis Atlas* (FMA).

### Comparison of Fish Metastatic Atlas (FMA) pages for cell lines with known differences in metastatic propensity

To test the capacity of our proposed pipeline to generate meaningful FMA pages we performed two positive control experiments: 1) We compared the pages for two metastatic melanoma cell lines with known stark differences in metastatic spread in mice [63]. Visually, the A375 cell line displayed broader spread along the vasculature (Fig. 4A). The MV3 cell line displayed a more granular pattern of discrete colonies. This may reflect the observation in mice where subcutaneous injection of the A375 cell line develops into widespread micrometastases, whereas the spread of the MV3 cell line is restricted to a single organ. Statistical comparison by permutation tests (see Methods, Supplementary Fig. S2) confirmed a significant difference (p<0.046) between the A375 and MV3 FMA pages. Interestingly, the difference in metastatic propensity between A375 and MV3 observed in mice could be predicted by an AI-based classifier that was trained on cell populations extracted from patient-derived melanoma xenografts (PDX), for which 1:1 correspondence between PDX behavior in the mouse and disease outcome in humans was established [64]. Thus, for this one example it seems plausible that the level of spread in the zebrafish over 48 hours relates to the metastatic disease progression in human patients. 2) We compared the pages for TC32 cells and NIH 3T3 fibroblasts (n>50). TC32 cells spread to the dorsal region, particularly the DLAV, which is only rarely occupied by the NIH 3T3 cells. In permutation tests these differences proved statistically significant (p<0.03; Fig. 4B). The broader occupancy of tissue by the TC32 cells may be the result of the genetic, epigenetic, and cell regulatory transformations these pediatric cancer cells have undergone. Thus, this finding readily shows the potential of the proposed pipeline to support molecular investigation of factors that will confine the metastatic potential of cancer cells.

**Fig. 4.**
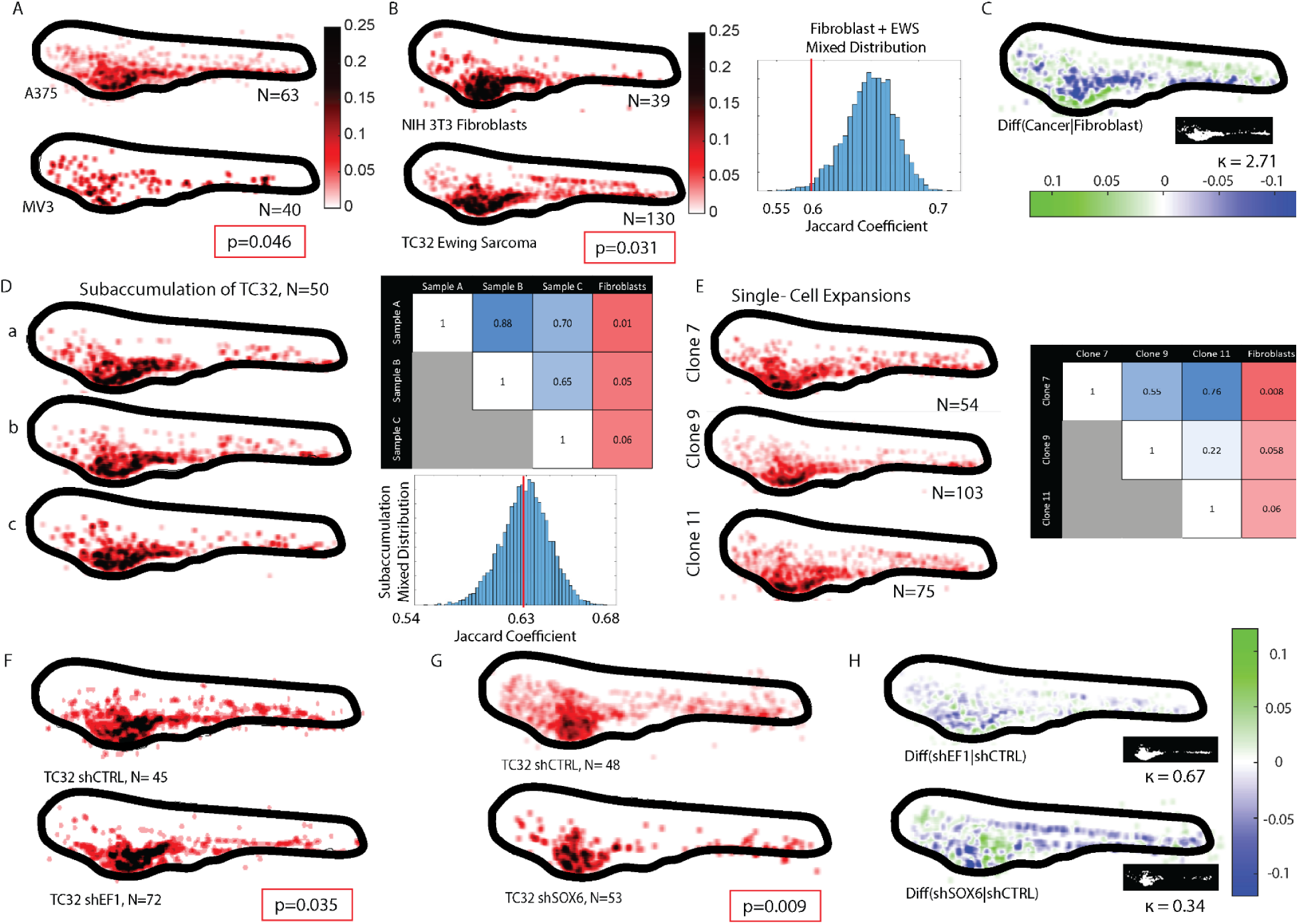
Differences in organotropsim in melanomas and in EwS cell lines with distinct oncogenotypes. **(A)** Accumulators for A375 and MV3 melanoma cell lines (classified in mice as highly and less metastatic, respectively). Box indicates P-value of a permutation test of the hypothesis that the two accumulators originate from the same sampling cohorts (see Methods). **(B)** Accumulators for NIH 3T3 fibroblasts and TC32 EwS cell line. Definition p-value, see (A). **(C)** Difference of accumulators in (B) using the fibroblast as a reference. Green colors indicate areas with higher probability of Ewing sarcoma colonies, blue colors areas with higher probability of fibroblast colonies. Inset, area of colony overlap (white). The value of k indicates the coefficient of accumulator divergence (see text). **(D)** Accumulators of 3 random sub-cohorts of N=50 embryos (sampled from a cohort of 109 embryos injected with the TC32 cell line). Discrimination matrix shows p-values of a permutation test of the hypothesis that the two compared accumulators originate from the same sampling cohort, which is confirmed by p-values >0.05. The last column of the matrix shows the p-values of comparing each TC32 sub-cohort accumulator the NIH 3T3 fibroblast accumulator in (B). **(E)** Accumulators of 3 single-cell clonal expansions of the TC32 cell line. Discrimination matrix shows p-values of a permutation test of the hypothesis that the two compared accumulators originate from the same sampling cohort, which is confirmed by p-values >0.05. The last column of the matrix shows the p-values of comparing each TC32 clone to the NIH 3T3 fibroblasts (B). **(F)** Accumulators for a TC32 cell line expressing scrambled shRNA (shCTRL) and a TC32 cell line expressing shRNA targeting the EWSR1:FLI1 fusion oncogene (shEF1). Definition P-value, see (A). **(G)** Accumulators for a TC32 cell line expressing scrambled shRNA (shCTRL) and a TC32 cell line expressing shRNA targeting the transcription factor SOX6 (shSOX6). Definition p-value, see (A). **(H)** Differences of accumulators in (F) and in (G) using the shCTRL-expressing cell lines as the reference. Insets, areas of colony overlap (white). k-values indicate the coefficient of accumulator divergence. **Note:** Most accumulators display outlier colonies beyond the outline of the template fish. They all tend to fall below the threshold of 2 visitations when generating masks for the computation of the Jaccard indices and thus, they have no influence on the statistical tests. Rather than removing them for visual presentations we retain these data points in the accumulators to document minor registration artifacts of individual larva especially in the chest region due to abnormal swelling after micro-injection. Outliers in the torso and trunk regions originate in slight registration mismatches for larva with large pitch, roll, or yaw. The registration filters in Fig. 2 have a false positive rate of ∼5%, causing some of these configurations to slip. conditions co-occupy. The coefficient is close to 0 for experiments in which the target condition minimally deviates from the co-occupied area and approximates infinity for experiments in which the reference minimally deviates from the co-occupied area. It is close to 1 when target and reference diverge approximately the same from the co-occupied area. The value κ = 2.72 indicates that the TC32 cancer cells spread substantially more throughout the fish than fibroblasts. In sum, these control experiments indicated to us the value of the FMA pipeline to quantify the spreading of human cancer xenotransplantations into the diverse tissues of a zebrafish embryo as a surrogate of metastatic propensity.

Because FMA pages directly show the probability of cells landing in a particular location, i.e. on a normalized scale, FMA pages registered to the same template can be readily subtracted to indicate areas of strong cell spreading differences (Fig. 4C). In addition, these difference maps allow the computation of a coefficient of distribution divergence between a target condition, in this example the TC32 cells, and a reference condition, in this example fibroblasts. The coefficient is defined as *k* = (*ρ*_*T*_ − *ρ*_*T*∩*R*_)⁄(*ρ*_*R*_ − *ρ*_*T*∩*R*_), i.e. the ratio of conditions co-occupy. The coefficient is close to 0 for experiments in which the target condition minimally deviates from the co-occupied area and approximates infinity for experiments in which the reference minimally deviates from the co-occupied area. It is close to 1 when target and reference diverge approximately the same from the co-occupied area. The value κ = 2.72 indicates that the TC32 cancer cells spread substantially more throughout the fish than fibroblasts. In sum, these control experiments indicated to us the value of the FMA pipeline to quantify the spreading of human cancer xenotransplantations into the diverse tissues of a zebrafish embryo as a surrogate of metastatic propensity.

### Organotropism of TC32 sarcoma xenografts likely originate from acute adaptation processes

We next wondered whether the significant heterogeneity between the TC32 spreading patterns between individual fish originated in variation of the clonal composition of the xenotransplants or reflected variation in the acute adaptation of cells to their microenvironments that would converge to a stable pattern of organotropism with a large enough number of repeat experiments. To address this question, we conducted two experiments: 1) We randomly selected subsamples of 50 fish from a total population of ∼550 fish injected with ∼100 TC32 cells. The differences between the spreading patterns for 3 representative samples were small by visual inspection and statistically insignificant by permutation tests. All 3 samples were significantly different from the fibroblast spreading pattern (Fig. 4D). 2) We expanded clonal subpopulations derived from single TC32 cells and generated FMA pages FMA pages for three randomly selected clones. Akin to random subsamples, the subclones were visually and statistically indistinguishable from one another, but all of them were different from the fibroblast pattern. Thus, we concluded that the heterogeneity in spreading patterns between individual fish does not originate in genomic variation of transplanted cells but is the outcome of variation in the acute adaptation of metastatic cells to the diverse microenvironments presented by the zebrafish.

### Shifts in organotropism indicate a role of the EWSR1::FLI1 oncogene as a promotor of cell plasticity

Previous work by us [65] and by others [66–68] has suggested that the fusion oncogene *EWSR1::FLI1* (EF1) may function as a plasticity factor. This would increase the aggressiveness of disease progression in general and metastatic spreading of EwS in particular by lending cells the ability to adapt broadly to different microenvironments. We therefore hypothesized that suppression of the expression of EWSR1::FLI1 would shift the spreading pattern and likely reduce the extension of tissue containing colonies. This hypothesis is countered by reports of EF1 knock-down promoting loss of cell-cell adhesion and acquisition of a motile, mesenchymal phenotype [67], which would yield a wider-spread distribution of metastatic colonies at the level of a whole organism. We generated FMA pages for TC32 cells expressing inducible scrambled control shRNA vs shRNA against EF1 gene expression (knock-down of the protein to <10%; see also Fig. S5a in Ref [65]). The spreading pattern of the two conditions were statistically different with p<0.035 (Fig. 4F). Visual comparison and accumulator difference maps (Fig. 4H, top) indicated a tendency of EF1 depletion to selectively reduce colonization of EwS cells in tissue surrounding the dorsal vasculature. However, the κ coefficient value of 0.67 suggested that this spreading defect was weaker than the difference between TC32 cells and fibroblasts.

Previous studies suggested that transformations of cells carrying the EF1 fusion gene are in part mediated by constitutively elevated expression of the transcription factor SOX6 [69]. Thus, we wondered whether knock-down of SOX6 would replicate the spreading deficiency of EF1 depleted cells (knock-down of the protein to 25 – 30%; see also Fig. S5 in Ref[69]). To our surprise, we found that SOX6 suppression reduced the TC32 spreading more pronouncedly than the knock-down of the upstream regulator EF1 (Fig. 4G, p <0.009, κ = 0.34). Similar to EF1 knockdown, SOX6 knockdown blocked colonization of tissues around the upper vasculature, in addition to lowering the spread to the head. While the stronger effect of SOX6 knockdown may simply the result of a differential stoichiometry in EF1-vs SOX6-mediated transcription, it is intriguing to point out the possibility that the functional penetrance of the EF1 fusion protein as an oncogene is weaker than the penetrance of some of its downstream targets owing to the multitude of pro- and anti-metastatic cell behaviors effected by EF1 transcriptional programs. The described pipeline now opens the door to systematic analyses of the contributions of EF1 targets to the organotropism of EwS.

### Patient-derived xenografts from primary tumors and lung metastasis show distinct patterns of organotropism

Finally, we sought to test whether FMAs would be able to reflect putative differences in organotropism between patient-derived EwS xenografts (PDXs). We contrasted PDXs derived from a pre-treatment biopsy of a paraspinal EwS and from a EwS relapse lung metastasis. Both tumor samples were passaged in mice to avoid artifacts accompanying cell culture (see Methods). Cells were harvested from primary tumors after dissociation of the tissue by trypsin and labelled by lentiviral transduction of a Tractin-mRuby reporter visualizing the actin cytoskeleton. Labelled cells were then injected in the PVS in the same quantities as with cell line xenografts. The same procedures of fish larvae selection and imaging were executed to assemble FMAs of the two PDXs. Both PDXs displayed substantial spreading (Fig. 5A), yet the accumulators were significantly different (p = 0.005, Fig 5B). Interestingly, the cells extracted from the lung metastasis occupied fewer sites (κ = 2.77, Fig. 5C). This could suggest that these cells have undergone a specialization, which makes them less agile for adaptation to diverse environmental conditions. While the presented data cannot rule out that the differences are driven by differences in the clonal composition of the two tumor types, our analysis of the equivalent spreading of different clones from the same cell population and of the effects of EF1 knock-down on organotropism leaves open the possibility that the lower apparent plasticity of the lung metastatic tumor results from the acquisition of tissue specific signaling circuits, which have been preserved throughout the xenografting in mice and fish. Future work exploiting the FMA pipeline as an indicator of tissue selectivity will be able to shed light on this fundamental question of pediatric cancer biology.

**Fig. 5.**
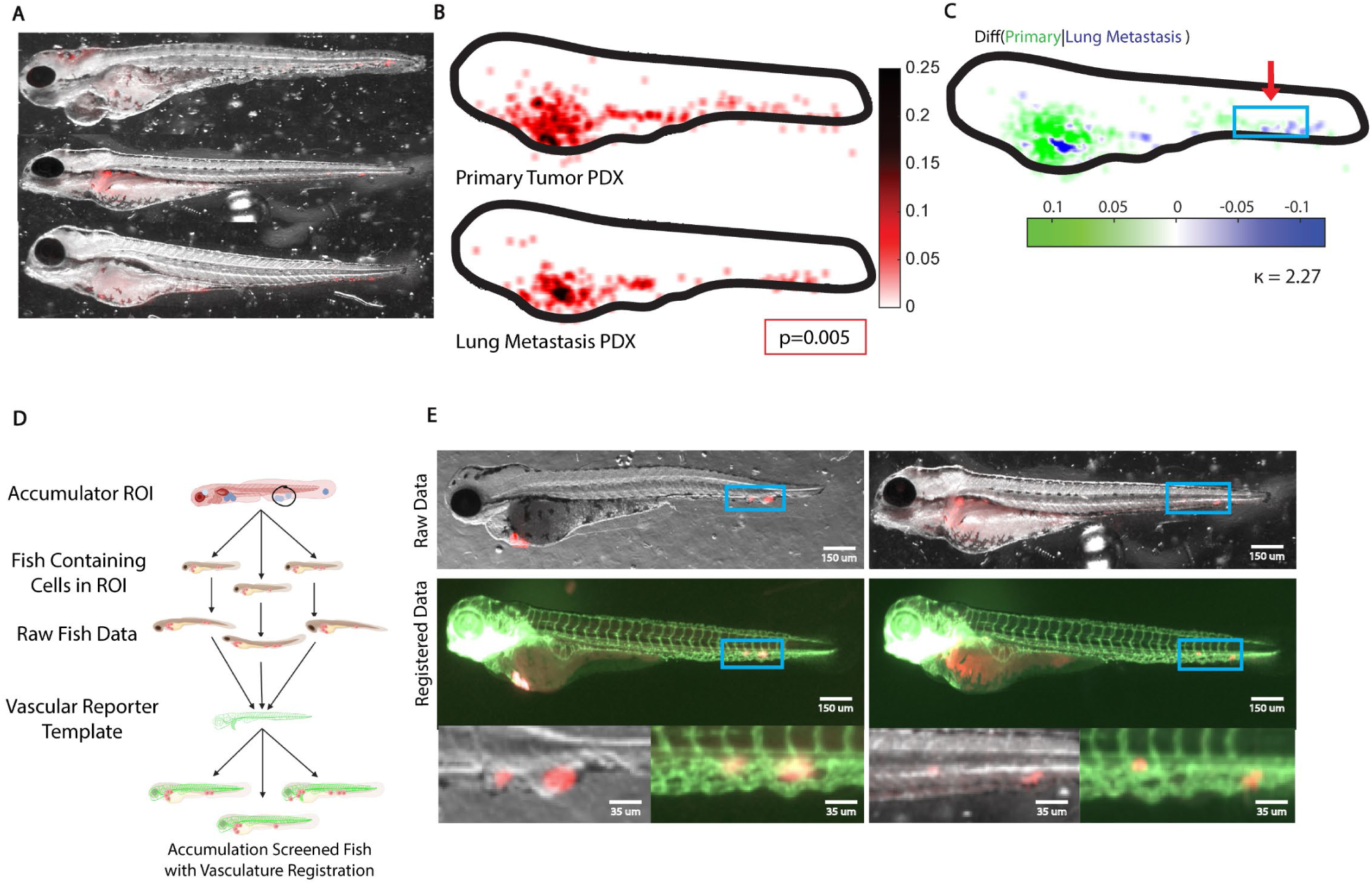
Organotropism of Ewing Sarcoma patient derived xenografts (PDX). **(A)** Injections of lentivirally labelled cells derived from a primary tumor. Shown are three representative fish embryos in brightfield images overlaid by the fluorescence signal of the xenografts. **(B)** Accumulators of PDXs of a primary tumor (top) and of a lung metastasis (bottom). **(C)** Differences of accumulators in (B) using the PDX from the lung metastasis as the reference. Green colors indicate areas with higher probability of PDX cells from the tumor to form colonies, blue colors areas with higher probability of PDX cells from lung metastasis to form colonies. κ-value indicates the coefficient of accumulator divergence. Box highlights a region of interest (ROI) with systematic difference in colony formation between the two PDXs. **(D)** Illustration of a computational workflow to identify embryos that contribute to the accumulator signal in the ROI, followed diffeomorphic mapping of the contributing colonies to the image of an embryo expressing vascular reporter for tissue context. **(E)** Two representative examples of embryos with colonies falling within the perimeter of the ROI indicated in (C) (top), their mapping onto a template of a Tg(fli:gfp) embryo (middle) and zoom ins of the ROI showing PDX colonies overlaid to the brightfield and vasculature reporter images of the template fish (bottom). **Note**: Outlier colonies beyond the outline of the template fish are visualized under the same considerations as discussed in Fig. 4.

The largest differences occurred in the ventral part of the tail (see Box, Fig. 5C). We wondered whether the differential would be associated with the posterior caudal vein. To test this, we made use of the bijectivity of registrations, both between the FMA page of the primary tumor spread and the individual constituent samples, and between the FMA page and a vascular reporter template (Fig. 5D): Given the selected region of interest (ROI), we first identified all constituent fish samples that contributed the signal of a metastatic colony to the metastatic hotspots in the ROI. Subsequently, we located these signals in the original raw fish images and computed bijective mappings from these images to the vascular reporter template. This allowed us to visualize the likely position of any of the detected colonies relative to the blood vessels. Fig. 5E shows representative examples of two colonies in the posterior caudal veins, the left originating from the primary tumor xenograft, the right from a from the lung metastatic tumor. Both colonies are localized just outside the blood vessels suggesting that these metastases are probably formed very close to the site of extravasation. In sum, these data illustrate the capacity of the proposed pipeline to detect differential organotropism between distinct PDXs and to associate the differential loci with specific host tissues in the zebrafish anatomy.

## Discussion

Our data outlines the operation of a pipeline for systematic and cost-effective screening of differential organotropism between distinct oncogenotypes or experimental conditions, harnessing the power of the zebrafish embryo as a host organism with the full diversity of tissue environment found in humans. Our work extends previous studies of metastatic propensity and drug sensitivity using zebrafish xenografts in its focus on differential colonization of host tissues [70–73]. We show sensitivity in distinguishing spreading patterns of; a pediatric sarcoma cell line vs an untransformed fibroblast; two melanoma cell lines with differential metastatic propensity as characterized previously in a mouse xenograft model of spontaneous metastasis; pediatric sarcomas in which we interfered with known oncogenic driver genes; and PDXs extracted from primary vs lung metastatic tumors. We supported these conclusions with negative control experiments, in which we showed that cell lines with identical oncogenotypes produce indistinguishable patterns of cell spreading. We also demonstrate that the pattern of cell spreading to physically and chemically diverse microenvironments by the EwS pediatric cancer model is not controlled via genetic selection, but likely arises from an elevated adaptability of this cancer to different environmental inputs. Almost certainly, this conclusion will not be universally applicable to all cancers. In fact, the proposed pipeline provides a powerful platform to identify cancers for which single cell clonal populations show distinct organotropism – a signature of genetic selection – versus cancers like the Ewing sarcoma model where single cell clonal populations show identical organotropism – a signature of non-genetic adaptation.

The main contribution of the proposed pipeline is the statistical rigor in distinguishing organotropism patterns in face of diverse sources of heterogeneity. Organotropism per se is the outcome of the heterogenous adaptation of cancer cells to diverse microenvironmental conditions. This biologically meaningful heterogeneity, which is at the core of cancer metastasis as a devastating disease is superimposed by experimental heterogeneity associated with anatomical variation between the hosts – here, zebrafish embryos – variation in sample handling and imaging, and meaningless biological and experimental noise. The technically critical steps to conquering the sources of meaningless heterogeneity include the acquisition of accumulator maps of cell spreading, referred to as pages of a fish metastasis atlas (FMA), from repeat sample sets large enough to suppress randomness while amplifying systematic behavior. In addition, we equipped the pipeline with simple statistical procedures to assess the significance of emerging patterns in FMA pages that may be indicative of differential organotropism between biological samples. To our knowledge this is the first implementation of a xenograft-based assay of organotropism of this level of experimental rigor and reproducibility.

### Limitations of the proposed pipeline

Despite the demonstrated potential uses for cancer research, at this stage the pipeline has several shortcomings that need to be factored into the conclusions and that warrant further technical refinements in future work:

1) As described here, the image analysis steps for embryo registration are specific to the imaging setup, in particular to the acquisition of a depth map that permits segmentation of the fins. While this has proven useful to constrain the fine-grained diffeomorphic mapping, easier alternatives may be available in cases where a second fluorescent channel can be spared for the alignment of the gross anatomy. Moreover, as illustrated in Fig. 2, a significant portion of the image analysis pipeline is concerned with the filtering of defects in the gross anatomy of the embryos and severe misalignments. This complication can be reduced by more efficient specimen handling such as enabled by a microfluidic platform like the VAST Bioimager[74]. Overall, there are potentially more convenient approaches to establishing diffeomorphism between accumulator templates and the individual larval specimens contributing to the FMA page of a certain oncogenotype/experimental condition.
2) The assay implicates several experimental variables that must be carefully controlled. Reproducibility at the level of the FMA pages depends on variables in the aquaculture of the fish itself, including overall health of the animals and feeding patterns, on the number of fertilized embryos, careful recalibration of the injection system with every session (see Methods) and on the injection site. To demonstrate the sensitivity of the spreading patterns on the injection site, we repeated the experiment with the MV3 cell model in Fig 4A using an injection of cells in the Duct of Couvier (DOC) instead of perivitelline space (PVS) (Fig. S3A). With injection into the DOC most cells are directly released into the blood stream. Thus, the spreading pattern reflects primarily the locations at which cells are able to extravasate from blood vessels and colonize secondary tissue. With PVS injections the spreading pattern also reflects the potency of the injected cells to invade the PV tissue and intravasate. As described in the section on xenotransplantation in our standard protocol we even remove fish with PVS injections that show cells immediately expelled into the vasculature. Thus, unsurprisingly the two injection protocols yield significantly different spreading patters (Fig. S3A). A further concern with PVS injections or with injections into other tissues from which cancer cells first have to actively enter the vasculature is that the strong accumulator signal around the injection site may unduly affect the Jaccard-based comparison of experimental conditions. To test this, we repeated the computation of p-values TC32 shCtrl and TC32 shEF1 with accumulators in which the yolk region was coarsely masked. The p-value of the comparison of the two EWS-FLI1 expression conditions changed marginal, without affecting the conclusion that EWS-FLI1 knockdown has a significant effect on organotropism (Fig. S3B). This robustness of the Jaccard-based statistics also dampens operator heterogeneity, both over time for one experimenter and between experimenters. The data acquired for this study relied on two experimenters, who were thoroughly cross trained. We verified the compatibility of their data using the bootstrap analyses as illustrated in Fig. 3, in which we assessed the statistical equivalence of intra- and inter-operator sessions. In sum, users of this assay are advised to select an injection site best compatible with the biological question and use only this site for a sensitive and robust comparison of different oncogenotypes and other experimental settings.
3) As implemented, the proposed assay generates merely an endpoint assessment of metastatic spreading that lumps all steps of the metastatic cascade into a single output pattern. The assay cannot distinguish niches that are colonized because they are easily accessible versus niches that are colonized because they allow elevated proliferation or survival. We believe that systematic differences in organotropism between oncogenotypes are more reflective of differences in the later steps of the metastatic cascade, i.e. the colonization, proliferation, and survival of metastatic niches and are likely less associated with differences between cancers in invasiveness, anoikis resistance in vasculature, and extravasation. Thus, we do not anticipate a particularly high accuracy of this assay in measuring metastatic propensity overall. This limitation is increased by the challenges in generating xenografts with reproducible initial mass and precise location. While these variations have minimal effects on the generation of particle-based accumulators across many fish, they will greatly complicate the analysis of intermediate stages in metastatic spreading.
4) While the zebrafish model offers numerous advantages for the analysis of organotropism, the level of predictive power for human disease progression remains unknown. As presented in Fig. 4A, we have tested two melanoma cell lines for which we have direct evidence of differential spreading in mice [63] and indirect evidence of differential metastatic propensity and disease outcomes in patients [64]. Their organotropic patterns in zebrafish align well with the expectations derived from these preclinical models and clinal observations. In Fig. 5 we also compare the spreading pattern of PDXs derived from a primary tumor and from lung metastases. The differential between the patterns has a plausible explanation in the context of human disease. Thus, a systematic relation of the organotropic behavior of human cancer cells in zebrafish and humans is conceivable. However, we are far from claiming that xenografts in the zebrafish will allow us to map 1-to-1 human cancer behavior. Our reservations originate primarily in the fact that with the proposed assay the effect of the host adaptive immune response to metastatic spreading is minimal. Even at longer time scales over which the zebrafish host mounts attacks of metastatic colonies by the immune system the accuracy of the pipeline to report the resistance of certain metastatic niches to immunosuppression for human cancers is unknown. The unique opportunities generated by the described work are 1) in providing a platform for relatively rapid and low cost analysis of the roles genetic and non-genetic factors play in driving metastatic spreading; 2) in unveiling the effects natural and genetically engineered environmental differences besides immune infiltration has on promoting differential response to chemotherapy and other stressors; 3) in offering a window to morphological and molecular underpinnings of such differential response via high-resolution imaging at multiple sites. Work is ongoing to employ FMA pages as a guide for single cell imaging as described in Segal et al. [65].

## Methods and Materials

### Zebrafish husbandry

*Danio rerio* were maintained according to industry standards in an AALAAC-accredited facility. AB and TL wild-type fish, and transgenic *fli:gfp* were obtained from the ZIRC Zebrafish International Resource Center (https://zebrafish.org) and were used for all xenograft injections.

### Zebrafish injections

Zebrafish were injected with human cell lines and PDX cells at 2 days post fertilization (dpf). The injection suspension nominally consists of 2×10^6^ cells (counted via Themo Fisher Countess) suspended in 20 µl PBS, corresponding to roughly 75-125 cells per fish applied over 1-2 injections. To account for variations in injection needle diameter and thereby microinjector pressure and time, the prepared cell suspension injections were counted in a drop of water on an agar-lined petri dish before every procedure. Pressure and injection time on the microinjector were adjusted accordingly to ensure this cell number was completed in 1-2 injections, while ensuring minimal stress to the embryo during injection (e.g. approaching excessive pressures or long injection times that depend on individual lab instrumentation). Injections were done at the perivitelline space between the yolk and ventral vasculature.

### Zebrafish xenograft screening and fixation

Zebrafish were screened under a Leica M205 FA fluorescent stereomicroscope 2 hours post injection (hpi) for cells in the tail, indicating potential non-biologically driven cell movement. These fish were removed. The remaining cell-positive fish were placed in a dark 34°C incubator and fixed at 48 hpi. Zebrafish were fixed in 4% PFA at 10°C overnight on a rocker.

### Cell labeling

All cell lines used for the generation of accumulators expressed fluorescent reporters in the RFP emission/excitation spectrum to allow for clearer imaging in fixed fish and minimize bleed-through of the high level of autofluorescence of fixed zebrafish larvae in the GFP channel. TC32 shSOX6 and their scrambled shRNA controls were labelled using a cytoplasmic mCherry fluorescent reporter. TC32 shEF1 and its scrambled shRNA control used a doxycycline inducible F-Tractin-mRuby reporter. The fibroblast (NIH3T3) cell lines, melanoma (A375, MV3) cell lines were labeled using a cytoplasmic mRuby reporter.

### PDX cell line generation

PDX cell lines (primary tumor, lung metastasis) were fluorescently labeled with a F-tractin mRuby reporter using the pLVX lentiviral system following standard protocols. Firstly, HEK293 cells were transfected with 5ug of each packaging vector and 5ug of the fluorescent plasmid using polyethylenimine (PEI). For every ug of DNA, 3ul of PEI was added. The mixture was added to HEK cells and following 24h incubation the media was collected and filtered through 0.45µm mixed cellulose esters membrane syringe filters (Fisher Scientific, Cat no. 09-720-005). The virus media and normal cell culture media were then added to the PDX cells in 1:1 ratio along with 2µg/mL polybrene (Millipore, Cat no. TR-1003-G). The cells were then selected using Neomycin (Geneticin Catalog number: 10131027).

### Imaging

Xenografted zebrafish were screened at 2 hpi for cells in the caudal vasculature away from the injection site. Fish were then fixed at 48 hpi, and immediately imaged on a Leica DM4000B fluorescent microscope operated through the LAS-X (Leica Software, 2019) software. During imaging, each fish was placed onto a glass coverslip with 34% Methylcelluose and oriented flatly on a sagittal plane. The microscope was equipped with a Leica DFC7000 T color CCD camera (Pixel size of 4.54×4.54 μm^2^) and a Leica Plan Apo 1x, M Series Common Objective Lens with variable NA, dependent on the magnification allowable for capturing in the entire larva in the field of view. For fish 48 hpi our magnification setting was between 3.60 and 4.02, translating into effective NAs of 0.07 and 0.08, respectively; and lateral resolutions of 4.5 μm and 4.1 μm, respectively. Given the camera pixel side length of 4.54 μm our lateral sampling rate was approximately 2x the Nyquist frequency. The range of effective NAs translated into depths of field between 85 and 100 μm. Hence, for sampling at the Nyquist frequency the allowable z-step in a Z-stack was between 42.5 μm and 50 μm. The acquisition of a Z-stack was prepared by manual focusing of the image at the top and at the bottom of a larva. The resulting vertical extension was then divided by the allowable z-step for the magnification, defining the number of slices necessary for capturing the entire larval volume (generally 12 – 20 slices). Z-stack images were then acquired in the brightfield (BF), autofluorescence (GFP), cell fluorescence (RFP) channels. Full-focus images were generated for each channel along with depth map image (DM) used for fish segmentation (see Main Text).

### Cell culture

TC32 shEF1, shCTRL, and shSOX6 cell lines described previously [69] (see Fig. 3A of this publication for SOX6 knock-down efficiency). EwS cell lines were fingerprinted via STR analysis and were confirmed multiple times during experimentation to be free of mycoplasma. All TC32 cells and shRNA constructs were maintained in (Gibco) RPMI 1640 GlutaMAX with 10% fetal bovine serum (Sigma) and 1× Antibiotic–Antimycotic (ThermoFisher) at 37°C in 5% CO_2_. A375 and MV3 cells were maintained in (Gibco) DMEM GlutaMAX with 10% FBS (Sigma) and 1× Antibiotic–Antimycotic (ThermoFisher) at 37°C in 5% CO_2_. PDX cells were cultured in DMEM GlutaMAX with 20% FBS (Sigma) at 37°C in 5% CO_2_.

All cell lines with the exception of PDXs were split or used for cell suspension using TrypLE Express (Gibco). PDX cells were split using Dispase in DMEM (Sigma-Aldrich SCM133). Because viability was a concern for culturing PDX cells, the gentler separation agent Dispase is highly recommended.

Cells containing inducible shRNAs via a tetracycline response element (TRE) were given doxycycline (Sigma Aldrich) at a concentration of 1µg/mL, replacing DOX every 48 hours. Cells were not injected until after 72 hours DOX exposure in culture to allow for a steady-state shRNA expression.

### Image Pre-processing and Segmentation

Xenograft full-focus images were checked and corrected via translation registration for any intra-stack drift between channels in the same fish resulting from micromovements in the 3% Methyl cellulose. Fish segmentation was performed as outlined in the Main Text and Fig. 1D.

### Image Registration

Segmented fish images were registered with a user-defined template fish by diffeomorphic mapping, i.e. for each pixel inside a target fish there is a unique location in the template fish and vice versa (see Main Text). The algorithm underlying the diffeomorphic mapping has been described and validated in Jiang et al. [60]

### Colony detection and Accumulation

Registered fish xenografted cell coordinates were detected by use of a Difference of Gaussian filtration (*σ*_1_ = 1, *σ*_2_ = 7 for cell lines and *σ*_1_ = 1, *σ*_2_ = 12 for the PDX samples) along with a global local maxima detection over the entire image. Maxima related to metastatic colonies were separated from non-specific maxima in the auto-fluorescence signal using Density Based Spatial Clustering Analysis for Applications with Noise (DBSCAN) [75], following the logic that cell data clusters are on average denser than random local maxima. The coordinates of cell clusters were copied into an accumulator, in which each cluster center was represented by a normalized unit Gaussian signal. The unit Gaussians from all contributing fish were summed in order to visualize areas of high cell visitation. To generate a final accumulator for an experimental condition, referred to as a Fish Metastatic Atlas (FMA) page, the raw accumulator was divided by the number of fish included in the analysis. Therefore, an FMA page reports in every pixel the probability of finding in any fish a metastatic colony on this location.

### Comparison of spreading patterns between different experimental conditions

To test whether two experimental conditions A and B produce systematically different spreading patterns we applied permutation testing of the Jaccard similarity coefficient:

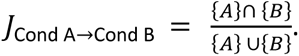

Here, {*A*} and {*B*} denote the sets of pixels in the accumulators of condition A and condition B, respectively, with a value above the user-defined visitation probability threshold (generally set to 2 colonies). The nominator thus describes the area of spreading overlap between the two conditions, and the denominator describes the area of the union of spreading.

The goal of the permutation test is to compute the probability *p*_*A*,*B*_ that the Jaccard coefficient *J*_Cond A→Cond B_ follows a distribution of overlaps between random spreading patterns. To sample this Null distribution we mixed the N_A_ and N_B_ images of detected colonies in individual fish into a joint pool of images (Suppl Fig. 2A). Two ‘null’ accumulators were then sampled pulling N_A_ and N_B_ random images from the joint pool. Both accumulators were binarized with the same probability threshold as the original data sets. The resulting Jaccard similarity coefficient produced one entry into the empirical null distribution (Suppl Fig. 2B). The procedure was then repeated *K* = 5000 times to construct. Given the null distribution of Jaccard coefficient values between randomly assembled accumulators the value for *p*_*A*,*B*_ is then computed as

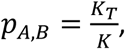

where KT is the number of elements in the null distribution with a value greater than *JJ*_Cond A→Cond B_ (Suppl Fig. 2C). All masks are generated at the same threshold as the permutation test.

We also assess a coefficient of distribution divergence between conditions A and B. The coefficient is defined as

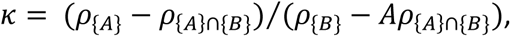

where *ρ*{.} denotes the area of a mask. The coefficient describes the ratio of differences in the spreading areas of the conditions *A* and *B* to the area both conditions co-occupy. The coefficient is close to 0 for experiments in which condition *A* minimally deviates from the co-occupied area and approximates infinity for experiments in which condition *B* minimally deviates from the co-occupied area. It is close to 1 when conditions *A* and *B* diverge approximately the same from the co-occupied area. We generally set condition *A* as the target condition and condition B as the reference, so that values *κ* >> 1 tend to indicate increased organotropism of the target and values *κ* << 1 tend to indicate reduced organotropism relative to a reference.

### Power Analysis of Xenografting Images

A statistical power analysis was done by comparing Jaccard similarity coefficients between two accumulators for probability thresholds of 1 and 2 visitations over an increasing number of fish images. Accumulators were generated by random sampling with replacement from 10 to 100 fish images per accumulator N=2000 times. These replicates were then averaged and plotted with their first derivative as a function of sample size.

To display the difference between a random point cloud distribution and a biological cell distribution, a singular Jaccard metric was calculated at probability thresholds of 1 and 2 visitations in random point cloud accumulators. These accumulators were generated by counting the total amount of cell events in the N=100 accumulator and randomly sampling that same amount with randomized positions inside of the template fish contour.

### PDX Implantation and Growth in vivo

Patient-derived xenografts (PDX) were generated at St. Jude Children’s Research Hospital and Nationwide Children’s Hospital, using previously described methods [76]. PDXs used in this study were EW-5 (derived from a pre-treatment biopsy of a paraspinal EwS) and NCH-EWS-1 (derived from a EwS relapse lung metastasis).

After passage in mice frozen PDX cells were transferred to the UT Southwestern labs, where they were thawed and orthotopically implanted in NOD-SCID mice as previously described [77]. Specifically, frozen cells were quickly thawed and added to 10 mL of warm DMEM in 15 mL conical, then spun for 10 minutes at 750 rpm. Supernatant was aspirated and cells were resuspended in 1:1 cold PBS to thawed Matrigel (Corning CLS356234) in 1 mL syringes then placed on ice. Female NOD-SCID mice were then subcutaneously injected with 100 µL Matrigel:cell suspension under sterile conditions. Animals were monitored twice weekly for tumor growth and were euthanized once tumors reached 2 cm^3^. This study was approved by the Institutional Review Board and the Institutional Animal Care and Use Committees of UTSW.

### PDX Tumor Dissociation

Harvested tumors were placed immediately in warm growth media (DMEM 10%FBS, 1% penicillin-streptomycin, 1% glutamine) for transport from animal facilities. Tumors were then finely minced using #11 sterile scalpel blades in 6 mm culture dish. Minced tissue was transferred to 50 mL conical containing a mixture of trypsin and collagenase IV (ThermoFisher 17104019) dissolved in DMEM and incubated at 37C for a minimum of 10 minutes until tissue easily fell apart on conical inversion. Dissociation process was then halted by adding STI, DNAse, and 1M MgCl and mixture was filtered using 40 µM strainer. Cell viability was determined using BioRad TC10 Automated Cell Counter and Trypan Blue stain. Cells were resuspended in PBS and proceeded to zebrafish embryo implantation

## Supplementary Figures

**Figure S1.**
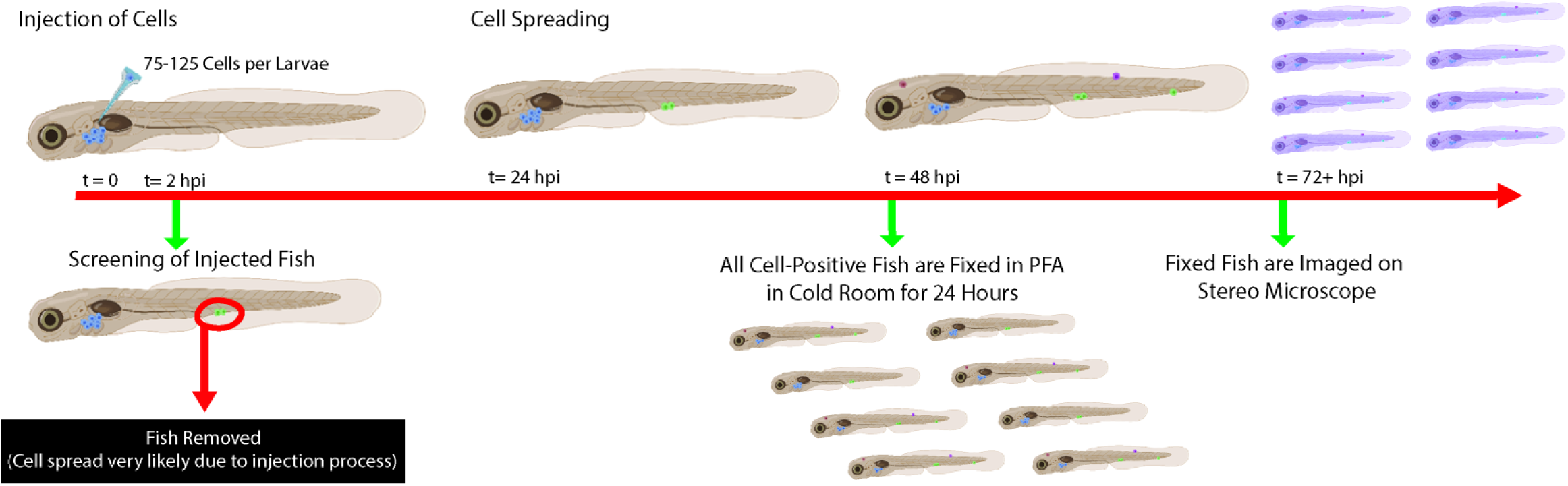
Cell injection workflow and processing timeline. Cells are injected at a concentration of 75-125 cells in 2-day old zebrafish larvae. Injected fish are visualized on a fluorescent stereoscope at 2 hours post injection (hpi). Fish containing cells in caudal vasculature are removed from the experimental population due to the high likelihood of cell deposition in these sites being non-biological and due to the injection process. Fish water changes (E3 media) are done every 18-24 hours along with removal of any dead larvae. Cell positive fish at 48 hpi are fixed in 4% PFA at 4°C overnight, washed and stored in PBS for imaging in fluorescent stereoscope or stored temporarily at 4°C.

**Figure S2.**
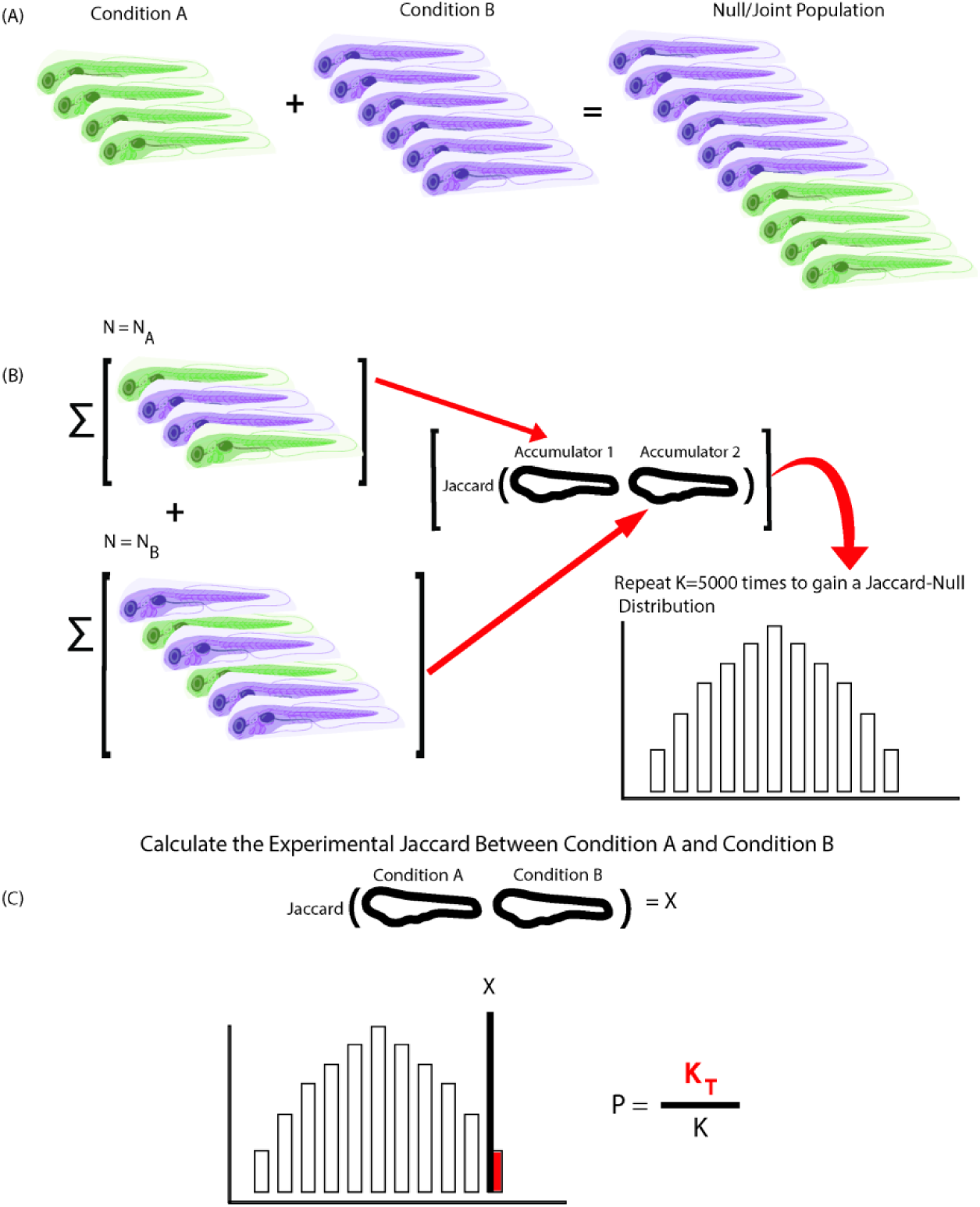
Statistical comparison of two Fish Metastatic Atlas pages. **(A)** Two fish cohorts of different experimental conditions A and B with a different number of fish (N_A_,N_B_) are mixed together to form a Null/joint population of N_A_+N_B_ fish. **(B)** N= N_A_ and N= N_B_ fish are randomly sampled (with no replacement) from the joint population in (A), followed by processing of two accumulators that are compared using the Jaccard similarity coefficient. The random sampling and comparison were repeated K=5000 times to generate a Jaccard Null-distribution for the joint data set. **(C)** The experimental Jaccard value X is computed between Condition A and Condition B accumulators. P-value is defined as the number of events in the Null-distribution less than X (K_T_) divided by the total number of randomized trials (K, K=5000.) **(D)** Example of randomly sampled Jaccard Null-distribution for the comparison of organotropic patterns for NIH 3T3 fibroblasts vs TC32 Ewing Sarcoma cell line. The experimental Jaccard value X is indicated in red, yielding a p-value of 0.031, i.e. the two patterns are deemed significantly different. See Fig. 4B. **(E)** Example of randomly sampled Jaccard Null-distribution for the comparison of organotropic patterns for sub-accumulation Sample A vs Sample B. The experimental Jaccard value X is indicated in red, yielding a p-value of 0.88, i.e. the two patterns are deemed statistically identical. See Fig. 4D. Note that the Null-distribution in (E) is narrower than in (D) because of the greater homogeneity of the cohort of fish making up the Null/joint population. This illustrates the need for specific sampling of Null-distributions with every comparison.

**Figure S3.**
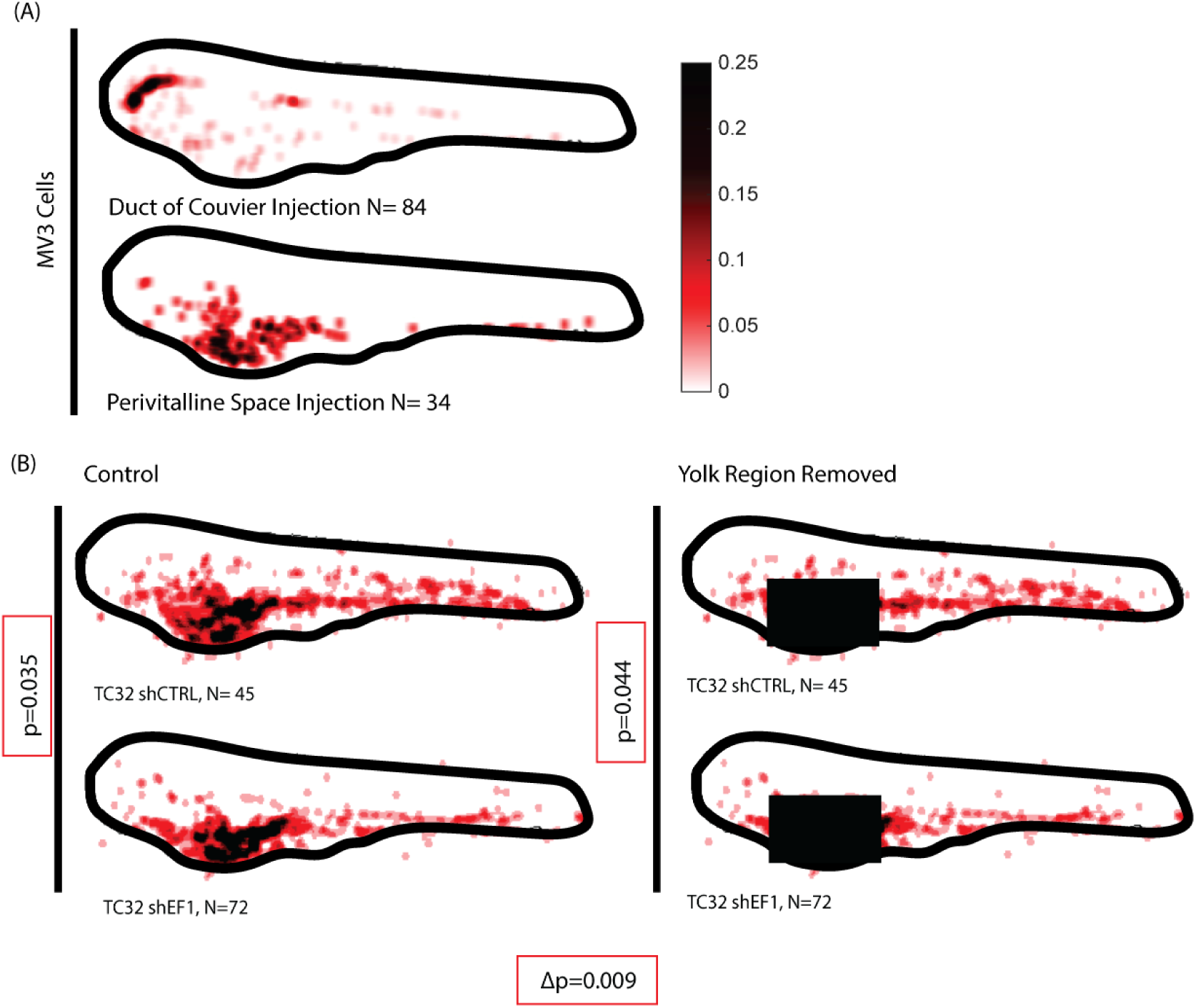
Effect of injection site choice and inclusion in the statistical comparison of Fish Metastatic Atlas pages. **(A)** Accumulators of MV3 xenotransplants into the Duct of Cuvier (top) and Perivitelline Space (bottom; this accumulator is similar to Fig. 4A). Box indicates P-value of a permutation test of the hypothesis that the two accumulators originate from the same sampling cohorts (see Methods). **(B)** Comparison of P-values of permutation tests of the hypothesis that TC32 expressing shCtrl or shEF1 (see Fig. Fig. 4F) generate the same sampling cohorts, once without (left) and once with the accumulator signal in the yolk mased. The difference in P-value between the two analyses is 0.009.

**Supplementary Video 1. Gross morphological differences between zebrafish larvae at 48 hours post injection.** Montage of N>100 fish to demonstrate the degree of variability in zebrafish larvae at the time of imaging. These examples are drawn from a large experimental cohort injected with TC32 EwS cells and allowed to grow for a further 48 hours before fixation and imaging.

## Acknowledgments

Funding for this study was provided by a grant from the National Cancer Institute (U54 CA268072 to J.A. and G.D.) and a fellowship (F31 CA236410 to D.S.). Additional funding in the Danuser lab was provided by the grant R35 GM136428, and in the Amatruda lab by grants U54CA231649 and P30CA014089. The Grünewald lab acknowledges funding by the Barbara and Wilfried Mohr foundation and the German Cancer Aid (DKH-111886 to T.G.P.G.). We thank Peter J. Houghton and the Childhood Solid Tumor Network (CSTN) at St. Jude Children’s Research Hospital for access to PDX models.

